# Reprogramming CAR T-Cells with designed bioPROTACs

**DOI:** 10.64898/2026.02.21.706835

**Authors:** Vivek S Peche, Sebastian Kenny, Tae Gun Kang, Brian Coventry, Tian Mi, Inna Goreshnik, Mariana Garcia Sanchez, Reid Martin, Macey Smith, Dionne Vafeados, Rahul S Kathayat, Yu Kaiwen, Zuo-Fei Yuan, Long Wu, Anthony High, Andrew Nemecek, Elizabeth Wickmann, Adeleye Adeshakin, Francesca Ferrara, Robert E Throm, Taosheng Chen, Benjamin Youngblood, David Baker, Stephen Gottschalk

## Abstract

Gene editing has been used to enhance CAR T-cell function by disrupting negative regulators but has limitations. Here we show that de novo-designed generated targeted degraders (bioPROTACs) provide an alternative approach. Expression of bioPROTACs in CAR T-cells targeting DNMT3A, a key regulator of T-cell exhaustion, phenocopied gene knockout. Our reversible, non-gene editing approach provides a tunable strategy to reprogram T-cell fate which should be broadly applicable for next-generation cell therapies.

## MAIN

Immunotherapy with CAR T-cells has shown great promise for hematological malignancies^1^. However, its antitumor efficacy for solid tumors is limited. One focus of enhancing CAR T-cell function is genome editing to disrupt genes that encode negative regulators^2^. For example, the DNA methyltransferase 3 alpha (DNMT3A) plays a critical role in orchestrating epigenetic program associated with T-cell exhaustion, and knockout of DNMT3A in CAR T-cells prevents exhaustion and enhances antitumor activity^3, 4^. However, DNMT3A and other genes that are currently targeted to enhance T-cell function are associated with pre-malignant states or clonal hematopoiesis of indeterminate potential^5, 6^. Thus, there is a need to develop approaches to transiently manipulate their function. While this could be achieved with small molecule inhibitors, these indiscriminately target all cells. To overcome these limitations, we set out to design proteins, biological proteolysis-targeting chimeras (bioPROTACs), to degrade DNMT3A and compare their effects to CRISPR/Cas9-mediated knockout (KO) in human CAR T-cells targeting the solid tumor antigen B7-H3. To design de novo proteins that specifically bind DNMT3A, we used RFdiffusion, a generative model that builds protein backbones matching a desired interface followed by ProteinMPNN for amino acid sequence design^7^ and AlphaFold2 for evaluating binding potential^8^ (**Fig. 1a**). From this pipeline, we identified 4,425 binders that passed computational structure prediction metrics. DNMT3A binding was validated by yeast surface display (**Fig. S1**) and 48 binders were selected for further development (**Fig. 1a,b**). These were fused to a synthetic E3 ligase-recruiting domain (hPEST degron or de novo Skp1 binder) to generate 96 DNMT3A degraders (D3ADegs). Plasmids encoding D3ADegs were transfected into a HiBiT-tagged DNMT3A reporter cell line (**Fig. S2a-c**) and four D3ADegs that decreased DNMT3A expression by >10% (range: 14.5-21.6%, **Fig. S2d**) were selected for further optimization (Online Methods section). A second screen identified two degraders (D3ADeg14, D3ADeg15) that decreased DNMT3A expression by >50% and ten by ≥20%, respectively (**Fig. S2e**). In the final screen only D3ADeg14 and D3ADeg15 decreased DNMT3A expression by >50% (**Fig. 1c**) without decreasing cell viability (**Fig. S2f**), and DNMT3A degradation was confirmed by western blot analysis (**Fig. S2g**). Based on these results, D3ADeg14 and D3ADeg15 were selected for further development. Both contained a FBXW11 domain and bound to the same region of DNMT3A but interacted with different amino acid residues (**Fig. 1d**).

**Figure 1:**
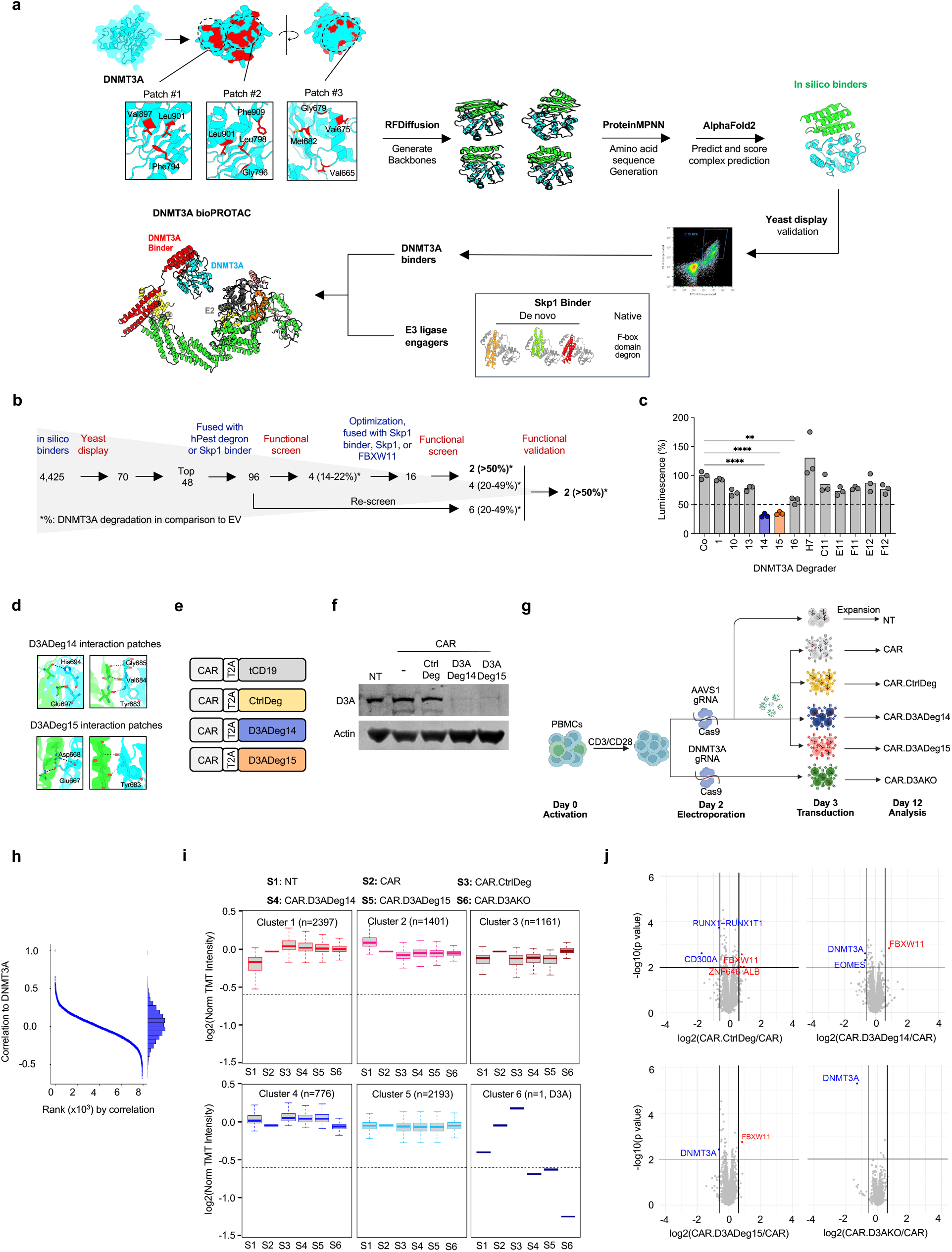
Generation and functional characterization of DNMT3A bioPROTACs. **a)** Workflow of DNMT3A binder design and screening. **b)** Summary of screening results. **c)** Functional validation screen of selected D3ADegs, two-way ANOVA with Dunnett’s multiple comparisons test, ****P <0.0001, ****P <0.01. **d)** Binding interface of D3ADeg14 and D3ADeg15. **e)** Scheme of retroviral constructs. **f)** Western blot of transduced T-cells. **g)** CAR T-cell generation scheme for comparison of CAR.D3ADeg14, CAR.D3ADeg15, and CAR.D3AKO T-cells. **h)** Protein correlation with DNMT3A, ranked by strength based on TMT quantitative proteomic analysis. **i)** Whole-proteomic weighted gene co-expression network analysis (WGCNA). **j)** Volcano plots of TMT proteomics for the indicated comparison.

To evaluate the ability of D3ADeg14 and D3ADeg15 to degrade DNMT3A in CAR T-cells we generated bicistronic retroviral vectors encoding a B7-H3-CAR and D3ADeg14, D3ADeg15, or control degrader (CtrlDeg) (**Fig. 1e**). Retroviral transduced T-cells demonstrated comparable CAR expression (**Fig. S3a**), and DNMT3A expression was only decreased in D3ADeg14 or D3ADeg15 T cells (**Fig. 1f**). There was no significant difference in CAR T-cell phenotype (**Fig. S3b-d**) and effector as judged by their cytolytic activity and ability to produce IFN-γ or IL-2 after exposure to B7-H3+ cells (**Fig. S3e-g**). To assess the biologic activity of the D3ADegs, we generated D3ADeg-expressing and DNMT3A KO (D3AKO) CAR T-cells (**Fig. 1g, S4a-c**) to perform Tandem Mass Tag (TMT) quantitative proteomics and functional analyses. Proteomic analysis showed broad and consistent peptide coverage, with DNMT3A emerging as the strongest self-correlated protein, confirming targeted degradation (**Fig. 1h**). Using weighted gene correlation network analysis (WGCNA), we identified six protein clusters, with DNMT3A presenting a distinct cluster (**Fig. 1i, Table S1**). Differential expression analysis showed significant reduction of DNMT3A in D3ADeg14, D3ADeg15 and D3AKO CAR T-cells, and as expected an increase of the FBXW11 degrader domain in D3ADeg CAR T-cells (**Fig. 1j**). These findings demonstrate that D3ADeg14 and D3ADeg15 specifically degrade DNMT3A.

To functionally compare D3ADeg14 and D3ADeg15 to D3KO, we performed a CAR T-cell repeat stimulation assay with tumor cells to mimic chronic antigen exposure (**Fig. 2a**). During repeat stimulation, we observed T-cell effector differentiation with preferential expansion of CD8+ CAR T-cells (**Fig. S4d**). CAR.D3ADeg14, CAR.D3ADeg15, and CAR.D3AKO T-cells exhibited sustained expansion, reaching significance for CAR.D3ADeg15 and CAR.D3AKO T-cells in comparison to control CAR T-cells (**Fig. 2b,c**). After the 1^st^ and 4^th^ stimulation we performed whole-genome bisulfite sequencing (WGBS) to compare the epigenetic landscape of CAR T-cells and used an established normalized multipotency index (MPI) that takes advantage of 245 CpG sites to predict their multipotent capacity^9^. After the 4^th^ stimulation, CAR.D3ADeg14, CAR.D3ADeg15, and CAR.D3AKO T-cells retained a high MPI, whereas CAR and CAR.CtrlDeg T-cells did not (**Fig. 2d**). Detailed methylation analyses after the 4^th^ stimulation revealed extensive DNA methylation changes between CAR and CAR.CtrlDeg versus CAR.D3ADeg14, CAR.D3ADeg15, and CAR.D3AKO T-cells (**Fig. 2e**). We identified 1,881 common differentially methylated regions (DMRs) in CAR.D3ADeg14, CAR.D3ADeg15, and CAR.D3AKO T-cells, which included genes highly associated with T-cell differentiation as judged by gene ontology (GO) analysis (**Fig. 2f,g**, **Table S2**) and two genes with well-established roles in T-cell stemness or longevity (*TCF7, LEF1*) are shown (**Fig. 2h**). A principal component analysis (PCA) using the post-4^th^ stimulation genome-wide DNA methylation profiles revealed that CAR.D3ADeg14, CAR.D3ADeg15, and CAR.D3AKO T-cells clustered together (**Fig. S5a**). Gene set enrichment analysis (GSEA) using the T-cell precursor exhaustion (Tpex) gene set, demonstrated enrichment in CAR or CAR.CtrlDeg T-cells in comparison to CAR.D3AKO or CAR.D3ADeg T-cells, respectively (**Fig. S5b**) with GO analyses shown for CAR.D3ADeg 14, CAR.D3ADeg15, and CAR.D3AKO T-cells (**Fig. S5c**).

**Figure 2:**
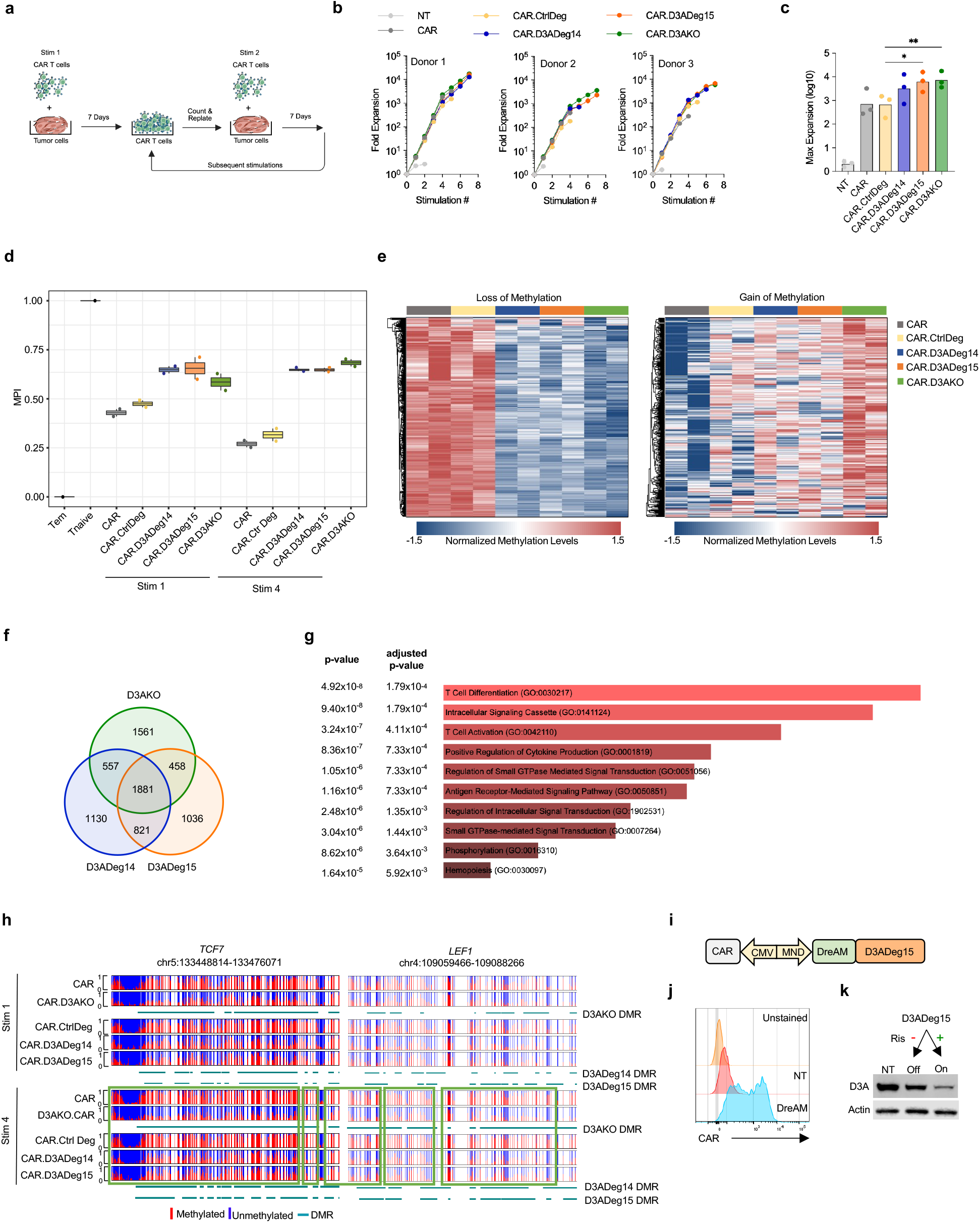
DNMT3A bioPROTACs preserve CAR T-cell plasticity. **a)** Scheme of the repeat stimulation co-culture assay. **b)** Expansion of CAR T-cells following weekly stimulations (n=3). **c)** Maximum (Max) expansion of CAR T-cells, two-way ANOVA with Tukey’s multiple comparisons test, *p <0.05, **p <0.01. **d)** MPI determined by WGBS after 1^st^ and 4^th^ stimulation (n=2). **e)** Heatmap showing top 300 differentially methylated and demethylated CpG sites after 4^th^ stimulation. **f)** Shared and non-shared DMRs between CAR.D3ADeg14, CAR.D3ADeg15, and CAR.D3AKO T-cells after 4^th^ stimulation. **g)** Gene Ontology (GO) analysis of shared DNMT3A-targeted genes with p-values, Fisher’s exact test. **h)** Representative methylation and demethylation events at *TCF7* and *LEF1* loci after 1^st^ and 4^th^ stimulation. Vertical lines: individual CpG sites (red: methylation, blue: no methylation). Green boxes: DMRs. **i)** Lentiviral construct with DreAM module. **j)** CAR expression of D3A 293T cells. **k)** Western blot of transduced and non-transduced D3A 293T cells +/-risdiplam (Ris).

To enable tunable degrader expression, we adapted a drug-elicitable alternative-splicing module (DreAM) approach that takes advantage of the FDA-approved drug risdiplam (**Fig. 2i**)^10^. Cells expressing DreAM-controlled D3ADeg15, only degraded DNMT3A in the presence of risdiplam (**Fig. 2j,k**). Thus, the DreAM is a readily translatable approach to control bioPROTACs in T-cells.

In summary, we have developed AI-based bioPROTACs to degrade DNMT3A in T-cells, which phenocopies DNMT3A KO. Unlike gene editing, which causes permanent deletion of the targeted gene, bioPROTAC expression can be regulated, which makes it ideal for developing programmable cell therapies. Our approach can be applied to any protein of interest and thus has broad applicability to adoptive cell therapies.

## Acknowledgements

We thank Chris DeRenzo for providing the retroviral vector encoding the B7-H3-CAR and Giedre Krenciute for the retroviral vector encoding DNMT3A-2A-CD20. Insertion-deletion (INDEL) analysis was performed by the Center for Advanced Genome Engineering (CAGE), which is supported in part by National Cancer Institute (NCI) P30 CA021765. This work was supported in part by the National Institutes for Health (NIH), including National Institute for Allergy and Infectious Diseases 1U19AI181881-01, National Institute on Aging R01AG063845, NCI R01CA240339 and Open Philanthropy Project Improving Protein Design Fund to DB, the American Cancer Society Postdoctoral Fellowship (PF-24-1318557-01-TBE) to SK, and the American Lebanese Syrian Associated Charities to BY and SG. The content is solely the responsibility of the authors and does not necessarily represent the official views of National Institutes of Health. Schematics shown in Figures 1g and 2a were created with BioRender (Biorender.com), for which we have a license.

## Authors contributions

Conceptualization: VSP, SK, DB, SG; Methodology: VSP, SK; Investigation: VSP, SK, TGK, BC, TM, IG, MGS, RM, MS, DV, RSK, YK, Z-FY, LW, AH, AN, EW, AA, FF, RT; Resources: VSP, SK,

DB, SG; Formal analysis: VSP, SK, TGK, TM, RSK, YK, Z-FY, AH, TC, BY, DB, SG; Supervision: DB, SG; Funding acquisition: BY, DB, SG; Writing – original draft preparation: VSP, SK, SG; Writing – review and editing: VSP, SK, TGK, BC, TM, IG, MGS, RM, MS, DV, RSK, YK, Z-FY, LW, AH, AN, EW, AA, FF, RT, TC, BY, DB, SG.

## REFERENCES

1. Cappell, K.M. & Kochenderfer, J.N. Long-term outcomes following CAR T cell therapy: what we know so far. Nat Rev Clin Oncol 20, 359–371 (2023).

2. Labanieh, L. & Mackall, C.L. CAR immune cells: design principles, resistance and the next generation. Nature 614, 635–648 (2023).

3. Ghoneim, H.E. et al. De Novo Epigenetic Programs Inhibit PD-1 Blockade-Mediated T Cell Rejuvenation. Cell 170, 142–157 e119 (2017).

4. Prinzing, B. et al. Deleting DNMT3A in CAR T cells prevents exhaustion and enhances antitumor activity. Science translational medicine 13, eabh0272 (2021).

5. Kang, T.G. et al. Epigenetic regulators of clonal hematopoiesis control CD8 T cell stemness during immunotherapy. Science 386, eadl4492 (2024).

6. Weeks, L.D. et al. Prediction of risk for myeloid malignancy in clonal hematopoiesis. NEJM Evid 2 (2023).

7. Cao, L. et al. Design of protein-binding proteins from the target structure alone. Nature 605, 551–560 (2022).

8. Jumper, J. et al. Highly accurate protein structure prediction with AlphaFold. Nature 596, 583–589 (2021).

9. Abdelsamed, H.A. et al. Beta cell-specific CD8(+) T cells maintain stem cell memory-associated epigenetic programs during type 1 diabetes. Nature immunology 21, 578–587 (2020).

10. Chen, Z. et al. The drug-elicitable alternative splicing module for tunable vector expression in the heart. Nat Cardiovasc Res 4, 938–955 (2025).

