## Supplemental Methods for "Reprogramming CAR T-Cells with designed bioPROTACs"

**In Silico Design of DNMT3A and Skp1 Binder**

The DNMT3A binders were generated using a protocol similar to the standard RFDiffusion protocol[1]. Protein Data Bank (PDB) 8TDR showcasing the methyltransferase domain of DNMT3A was used as the targeting structure. Three hydrophobic hotspot patches were chosen to design against. The three patches and the hotspot residues chosen are shown in detail in **Fig. 1A**. 10,000 diffusion trajectories designing 80-120 amino-acid binders were produced and the structures from several early timesteps were output in addition to the final structure. These outputs were designed with ProteinMPNN, filtered by Rosetta ddG, and 300,000 were predicted with AlphaFold2 initial guess. The best 5,000 structures passed the scoring metric of predicted local distance difference test (pLDDT) > 90, in addition to a combination of Rosetta ddG and AlphaFold2 pae_interaction were ordered as gene fragments and tested by yeast surface display. Skp1 binders were designed for E3 ligase recruitment using PDB 1LDJ as the targeting structure. Hydrophobic residues on the F-box interface were used as hotspots. Similar filtering methods were used to choose designs to order for yeast surface display screening.

**Yeast Surface Display**

The DNMT3A binders and Skp1 binders generated were ordered as gene fragments (Twist Biosciences) and cloned into pETcon vector, which encodes a c-Myc tag, for yeast expression. The cloned plasmid was transformed into EBY100 yeast strain using a previously established protocol[2]. Transformed yeast cultures were grown in C-Trp-Ura medium supplemented with 2% (w/v) glucose. For induction of expression, yeast cells were grown in SGCAA medium supplemented with 0.2% (w/v) glucose at the cell density of $1 \times 10^7$ cells per mL and induced at 30°C overnight. For DNMT3A binder screening, Cells were washed with PBSF (PBS with 1% (w/v) BSA), incubated with commercially available DYKDDDDK(FLAG)-DNMT3A (Active Motif, Cat #31406), washed and secondarily stained with PE anti-DYKDDDDK (BioLegend, Cat #637309) and anti-c-Myc fluorescein isothiocyanate (Miltenyi Biotech, Cat # 130-116-485). For Skp1 binder screening, cells were washed with PBSF (PBS with 1% (w/v) BSA), incubated with commercially available biotinylated Skp1 (Sino Biological, Cat # 14161-H40E-B), washed and secondarily labeled with Streptavidin R-Phycoerythrin Conjugate (Thermo Fisher, Cat #S866) and anti-c-Myc fluorescein isothiocyanate (Miltenyi Biotech, Cat # 130-116-485). Two rounds of sorts on the Sony SH800 Cell Sorter with different concentrations of FLAG-tagged or biotinylated targets were performed to enrich strong binders before a final round of titration sort to deduce the 50% saturation concentration ($SC_{50}$) of the enriched DNMT3A binders.

### Design and Cloning of DNMT3A bioPROTAC Library

The verified DNMT3A binders were ordered as gene fragments (Twist Biosciences) and cloned into an in-house CMV-based vector compatible with golden gate assembly. The fragments ordered included DNMT3A binders fused to C-terminus of the human (h)PEST degron sequence[3,4], Skp1 binder fused to DNMT3A binders, and a fragment of human FBXW11 (amino acid 33 to 228) containing the F-box domain fused to DNMT3A binders. All fusions were performed using 10xGly-Ser linkers. The cloned plasmids were transformed into DH5α strain (NEB, Cat #C2897H) and inoculated in Terrific Broth II supplemented with ampicillin for amplification. The amplified plasmids were extracted from the cultures by miniprep in either a 96-well format for screening as described in the Screening of DNMT3A bioPROTACs section or individually (Qiagen, Cat #27106) for large-scale transfections.

### Cell Lines

The A549 lung adenocarcinoma and HEK293T human embryonic kidney cell lines were obtained from the American Type Culture Collection (ATCC,USA). The osteosarcoma cell line LM7 was kindly provided by Eugenie Kleinerman (The University of Texas MD Anderson Cancer Center, Houston, Texas, USA) as previously reported[5]. All adherent tumor cell lines were maintained in Dulbecco's Modified Eagle Medium (DMEM, Gibco) supplemented with 10% fetal bovine serum (FBS, Gibco) and 1% GlutaMAX (Gibco), and were subcultured using 0.05% trypsin-EDTA (Thermo Fisher Scientific). HEK293T cells expressing DNMT3A (D3A 293T reporter cells) were generated by standard retroviral transduction with pSFG-CD20.HiBiTD3AFlag. Cell lines were authenticated using ATCC's human STR profiling authentication service. Cell lines were routinely checked for mycoplasma by the MycoAlert Mycoplasma Detection Kit (Lonza) and were negative.

### Generation of Retroviral Vectors

The following retroviral constructs were used in this study: pSFG-B7-H3-CAR.tCD19 (CAR), pSFG-B7-H3-CAR.CtrlDeg (CAR.CtrlDeg), pSFG-B7-H3-CAR.D3ADeg14 (CAR.D3ADeg14), pSFG-B7-H3-CAR.D3ADeg15 (CAR.D3ADeg15), and pSFG-CD20.HiBiTD3AFlag. The B7-H3-CAR construct was previously published and consisted of an antigen binding domain derived from the monoclonal antibody (mAb) MGA271, a CD8α hinge and transmembrane domain, and a CD28.ζ signaling domain[6,7]. The CAR retroviral vector was generated by inserting a self-cleaving T2A-tCD19 sequence into the C-terminus of the pSFG-B7-H3-CAR.CD28.ζ vector using In-Fusion cloning (Takara Bio) according to the manufacturer's protocol. Using the CAR retroviral vector as the backbone, constructs encoding the control degrader, DNMT3A Degrader 14

(D3ADeg14), and DNMT3A Degrader 15 (D3ADeg15) were similarly cloned downstream of the T2A sequence by In-Fusion cloning. The pSFG-CD20.HiBiTD3AFlag vector was generated by inserting an N-terminal HiBiT tag into the previously published pSFG-CD20.D3AFlag vector using In-Fusion cloning[8]. All DNA oligonucleotides for vector amplification and inserts (including degrader sequences, HiBiT tags, and linkers) were synthesized as double-stranded gene fragments (gBlocks) and ordered from Integrated DNA Technologies (IDT). All generated constructs were verified by sequencing (Plasmidsaurus).

RD114-pseudotyped retroviral particles encoding CAR, CAR.CtrlDeg, CAR.D3ADeg14 and CAR.D3ADeg15 were produced by transient transfection of HEK293T cells as previously described[8]. Viral supernatants were harvested at 48 hours post-transfection, centrifuged at 400xg for 5 minutes (min) at 4°C to remove cell debris, filtered through a 0.45 μm PES filter, and either used immediately for same-day T cell transduction or snap-frozen in 2-3 mL aliquots and stored at -80 °C.

**Generation of HiBiT DNMT3A Reporter Cell Line**

HEK293T cells were transduced with retroviral particles encoding pSFG-CD20.HiBiTD3AFlag. For transduction, 24-well tissue culture-treated plates were coated with Retronectin (Takara Bio, 10 μg/mL in PBS) and incubated overnight at 4°C. Plates were washed once with PBS, followed by addition of 500 μL of retroviral supernatant per well. Plates were centrifuged at 2000xg for 90 min and $2.5 \times 10^5$ HEK293T cells in 500 μL of DMEM (Gibco) supplemented with 10% FBS (Gibco) were added per well. Cells were incubated overnight at 37 °C in a humidified incubator with 5% $CO_2$. The following day, viral supernatant was replaced with fresh DMEM supplemented with 10% FBS. Cells were expanded and HiBiT-DNMT3A expression was confirmed by Nano-Glo HiBiT Lytic Assay (Promega) and Western blot using anti-DNMT3A (clone D23G1, Cell Signaling Technology); anti-GAPDH (clone 0411, Santa Cruz Biotechnology) was used as loading control.

**Screening of DNMT3A bioPROTACs**

Screening was performed using a reverse transfection of the HiBiT DNMT3A reporter cell line in white, flat-bottom 384-well plates (Revvity, 6007688). D3ADegs encoding plasmids or empty vector control were complexed with either Lipofectamine 3000 (Invitrogen) or FuGENE6 transfection reagent (Promega) in OptiMEM (Gibco). For Lipofectamine 3000 based transfections, 29 ng of each plasmid were dispensed to 384-plate using acoustic liquid handlers (Beckman Coulter, Labcyte Echo 655T). 10 μL of transfection master mix containing OptiMEM (9.8 μL),

P3000 (0.1 µL) and Lipofectamine 3000 (0.075 µL) were added to each well, plate was centrifuged at 1,200 rpm for 1 minute (min), shaken on Multidrop Combi (Thermo Scientific) for 30 seconds (s), and centrifuged again at 1,200 rpm for 1 min. For FuGENE6 based transfections, 75 ng of each plasmid were dispensed to 384-plate using Echo 655T. 9.3 µL of transfection master mix containing OptiMEM (8.9 µL) and Fugene 6 (0.4 µL) were added to each well, plate was centrifuged at 1,500 rpm for 1 min and then shaken on Multidrop Combi for 1 min. After a 15 min incubation at room temperature, 7K, 7.5K, or 10K cells per well were seeded in15 µL of DMEM supplemented with 1x P/S and 10% fetal bovine serum using an automated liquid dispenser (Thermo Scientific, Multidrop Combi 836). Plate was shaken on Multidrop Combi for 1 min, centrifuged at 500 rpm for 1min and incubated at 37 $^{o}$C. HiBiT assay was performed at 24h and 48h post-transfection using the Nano-Glo HiBiT Lytic Detection System (Promega, N3050). Briefly, at each time point, plates were equilibrated to room temperature for 10 min. 25 µL of lysis buffer containing LgBiT and substrate was dispensed into each well using Multidrop Combi and resulting plate was shaken for 30 s. The plate was centrifuged at 1000 rpm for 1 min, incubated at room temperature for 20 min, and followed by luminescence measurement on microplate reader (PerkinElmer, EnVision 2104). Raw luminescence values for each bioPROTAC were normalized to the mean of empty vector control wells on each plate. For selected bioPROTACs, cell viability was determined using a CellTiter-Glo (CTG) assay (Promega, G7570).

**Generation of B7-H3 CAR T-Cells**

Peripheral blood mononuclear cells (PBMCs) were isolated from purchased leukapheresis products (Charles River) by density gradient centrifugation using Lymphoprep (STEMCELL Technologies) and subsequently cryopreserved. St Jude Children's Research Hospital Institutional Review Board reviewed the research activity and determined it to be non-human subject research since the leukapheresis products were de-identified. For generation of non- and gene-edited CAR T-cell populations we followed previously published protocols of our laboratory[8].

For initial stimulation, PBMCs were plated at $1x10^6$ cells per well in 24-well tissue culture-treated plates pre-coated overnight at 4°C with 0.5 mL of sterile water containing 250 ng each of anti-CD3 (Miltenyi Biotec, #130-093-387) and anti-CD28 (Miltenyi Biotec, #130-093-375) monoclonal antibodies. Cells were cultured in RPMI 1640 supplemented with 10% heat-inactivated fetal bovine serum (FBS) and 2 mM GlutaMAX (Thermo Fisher Scientific). Twenty-four hours after activation, recombinant human IL-7 (10 ng/mL) and IL-15 (5 ng/mL) (PeproTech) were added to support proliferation and survival.

On day 2, genome editing was performed by electroporation of Cas9-sgRNA ribonucleoprotein (RNP) complexes with validated guides targeting DNMT3A or AAVS1 (control)[8]. RNPs were prepared by mixing 3 µL of 60 µM sgRNA (IDT) with 1 µL of 40 µM Cas9 (MacroLab, UC Berkeley) and incubating for 10 minutes at room temperature. Complexes were stored at -80°C until use. For single-gene knockout, $6x10^5$ activated T cells were resuspended in 20 µL of Neon R buffer and mixed with 4 µL of the appropriate RNP. Electroporation was performed using the Neon Transfection System (Thermo Fisher Scientific) with three 10 ms pulses at 1,600 V. Two 10 µL electroporated reactions were pooled and transferred to 48-well tissue culture–treated plates containing 500 µL of prewarmed RPMI 1640 with 20% FBS, 2 mM GlutaMAX, IL-7 (10 ng/mL), and IL-15 (5 ng/mL).

On day 3, cells were transduced in a non-tissue culture-treated 24-well plates that were coated the night before with Retronectin (5 µg/well in 500 µL PBS) and stored at 4°C. Plates were washed once with PBS, and loaded with 500 µL/well of RD114-pseudotyped retroviral supernatant encoding CAR, CAR.CtrlDeg, CAR.D3ADeg14 or CAR.D3ADeg15. Viral plates were spinoculated at 2000xg for 90 min. T-cells were resuspended at $1.25\,x10^5$ cells/mL in RPMI with IL-7 and IL-15, and 2 mL ($2.5\,x10^5$ cells) was added per well. Transduced T cells were maintained in cytokine-containing media and transferred to tissue culture-treated plates on Day 5 or 6. Cytokines were replenished every 2-3 days. Cells were used for downstream analyses or cryopreserved between Days 7 and 14 post-transduction. Editing efficacy was confirmed by INDEL analysis.

For experiments with non-edited B7-H3-CAR T cells the same protocol was followed with omission of the gene-editing step and activated T cells were transduced on Day 2.

**Western Blot**

CAR T-cell population were harvested on day 7 post-transduction, washed once with PBS and lysed in RIPA buffer (Cell Signaling Technologies) supplemented with Halt Protease and Phosphatase Inhibitor Cocktail (Thermo Fisher Scientific). Protein concentration was determined using the BCA protein assay (Thermo Fisher Scientific). Equal amounts of total protein (30 µg) were resolved by SDS-PAGE and transferred to nitrocellulose membranes (Bio-Rad) using standard wet transfer protocols. Membranes were blocked for 1 hour at room temperature in LI-COR blocking buffer and incubated overnight at 4°C with the following primary antibodies: rabbit anti-human DNMT3A (clone D23G1, Cell Signaling Technologies), mouse anti-human GAPDH (clone 0411, Santa Cruz Biotechnology), or HRP-conjugated mouse anti-actin (clone C4, sc-

47778, Santa Cruz Biotechnology). After 3x washing with TBS-T, membranes probed for DNMT3A or GAPDH were incubated for 1 hour at room temperature with IRDye-conjugated secondary antibodies (goat anti-rabbit IRDye 680CW or goat anti-mouse IRDye 800RD, LI-COR). Blots probed for actin were developed using SuperSignal West Femto Maximum Sensitivity Substrate (Thermo Fisher Scientific) and imaged by chemiluminescence. LI-COR Odyssey imaging was used for IRDye detection and chemiluminescence.

**Co-Culture Assay**

CAR T cells or nontransduced T cells were co-cultured with B7-H3$^{+}$ LM7 tumor cells[6] at a 2:1 effector-to-target (E:T) ratio in 24-well plates. After 24 hours of incubation, culture supernatants were collected, and the concentration of IFN-γ and IL-2 was measured using ELISA according to the manufacturer's protocol (Ella Automated Immunoassay System, Bio-Techne Corporation).

**Real-Time Cytotoxicity Assay Using xCELLigence RTCA**

The cytolytic activity of CAR T cells against tumor cells was assessed using the xCELLigence Real-Time Cell Analysis (RTCA) DP instrument (Agilent Technologies). LM7 target cells were seeded in E-Plate 96 (Agilent) at a density of $4 \times 10^4$ cells/well in complete RPMI media and incubated overnight until a stable baseline Cell Index was established. The next day, effector CAR T cells (CAR, CAR.CtrlDeg, CAR.D3ADeg14, CAR.D3ADeg15) cells or nontransduced T cells were added at a 2:1 effector-to-target (E:T) ratio. Impedance, expressed as Cell Index, was recorded every 15 minutes for 48 hours using RTCA software according to the manufacturer's instructions. Cytotoxicity was calculated by comparing the reduction in Cell Index to that of target cell-only control wells.

**Repeat Stimulation Assay**

Control KO (Ctrl), DNMT3A KO, Ctrl Degrader and DNMT3A degrader (Deg14 or Deg15) CAR T-cells were cocultured with A549 tumor cells at an effector-to-target (E:T) ratio of 2:1 in complete RPMI medium supplemented with IL-15 (5 ng/mL) as previously described[8]. After 7 days, T-cells were harvested, counted, and restimulated with freshly plated A549 cells at the same E:T ratio in the continued presence of IL-15. This stimulation cycle was repeated weekly. At each round, T-cells were monitored for expansion and cytolytic activity. Stimulation continued until T cells no longer demonstrated measurable tumor killing.

**Drug-Elicitable Alternative-Splicing Module (DreAM) Expression System**

To demonstrate tunable expression of bioPROTACs, we inserted into a bidirectional MND/minimal CMV promoter expression cassette into a self-inactivating (SIN) 3′ partially deleted viral long terminal repeats (LTRs) lentiviral vector (LV). The B7-H3-CAR.CD28.ζ was cloned downstream of the minimal CMV promoter and the Drug-Elicitable Alternative-Splicing Module (DreAM) sequence responsive to risdiplam[9] was cloned downstream of MND promoter followed by the coding sequence of D3ADeg15 in which the six methionines in FBXW11 were replaced with leucines (rev.B7-H3-CAR.DreAM.D3ADeg15). VSVG-pseudotyped LV retroviral particles were produced by transient transfection and used to transduce the D3A 293T reporter cell line. Following transduction, cells were treated with 10 µM risdiplam (Tocris Bioscience) for 48 hours and processed for Western blot analysis.

**Bisulfite Sequencing Methylation Profiling and Data Analysis**

DNA extraction and enzymatic conversion were performed using NEBNext Enzymatic Methyl-seq Kit. Converted genomic DNA were submitted to the Hartwell Center at St. Jude Children's Research Hospital for library construction and sequencing on the Illumina NovaSeq platform with 150bp paired-end and target reads of 500 million per sample. EM Seq reads were trimmed by 1bp on 5' and 3' ends as well as all Illumina adapter sequences. Trimmed reads were aligned to the hg19 genome using the BSMAP v. 2.90 software. Methylation levels were called by methratio.py script of BSMAP. Differential methylation analysis was performed by R package DSS 2.34. Basic two group comparisons were analyzed using a cutoff of $p \leq 0.01$. T cell multi potency index was calculated based on 245 CpGs and weights as published[10]. DMRs with more than 20% methylation differences were counted in Venn Diagram. And top 300 DMRs were plotted in heatmaps and used as input for Enrichr GO analysis (https://maayanlab.cloud/Enrichr/). Gene set used for GSEA analysis was generated from a published supplementary table[11]. Pre-ranked GSEA was ran on DMR tables with -sign(logFC)*log10(areaStats) of most significant DMR of a gene as the rank.

**Immunophenotyping by Flow Cytometry**

For immunophenotypic analysis, CAR T cells were harvested, washed with PBS, and stained in FACS buffer (PBS with 2% FBS) for 20 min at 4°C in the dark. Transduced cells were detected by anti-G4S (Cell Signaling Technology, 69782S) or anti-human F(ab'2) antibody (Jackson ImmunoResearch Laboratories, Inc., 109-606-006). The following fluorochrome-conjugated monoclonal antibodies (all from BD Biosciences) were used to assess T-cell subsets and memory phenotype: BV421 anti-human CD3 (clone UCHT1, Cat# 563796), PE-Cy7 anti-human CD4 (clone SK3, Cat# 557852), APC-H7 anti-human CD8 (clone SK1, Cat# 560179), FITC anti-human CCR7 (CD197) (clone 150503, Cat# 561271), PerCP-Cy5.5 anti-human CD45RA (clone HI100, Cat# 563429), LIVE/DEAD™ Fixable Aqua Dead Cell Stain (ThermoFisher Scientific eBioscience™ Cat. No. L34957). Following surface staining, cells were washed and resuspended in FACS buffer and analyzed on a BD LSRFortessa flow cytometer. Data were compensated and analyzed using FlowJo software (BD Biosciences). T cell subsets were gated on viable singlet $CD3^+$ lymphocytes, and memory phenotype was determined based on CCR7 and CD45RA expression.

**Tandem-Mass-Tag (TMT) Labeling and Two-Dimensional Liquid Chromatography-Tandem Mass Spectrometry (LC/LC-MS/MS)**

CAR, CAR.CtrlDeg, CAR.D3ADeg14, CAR.D3ADeg15, D3AKO.CAR T cells were homogenized in the lysis buffer (50 mM [4-(2-hydroxyethyl)-1-piperazineethanesulfonic acid] (HEPES), pH 8.5, 2% sodium dodecyl sulfate (SDS), 1x PhosSTOP phosphatase inhibitor cocktail (Roche), and 5 mM dithiothreitol (DTT)). The proteins were alkylated with 10 mM iodoacetimide (IAA) for 30 min in the dark, and quenched with 10 mM DTT for 20 min, all at room temperature (RT), then subjected to paramagnetic beads cleanup and trypsin digestion as described[12]. Briefly, a 1:1 mix of two types (hydrophilic (Sera-Mag Speed Beads, GE Life Sciences, CAT# 45152105050250) and hydrophobic (Sera-Mag Speed Beads, GE Life Sciences, CAT# 65152105050250)) of beads were added to proteins at a bead to protein ratio of 5:1 (w/w). 100% ethanol was added to the mixture to reach a final concentration of 50%, and then incubated on a shaker for 10 min at 1000 rpm at RT. The supernatant was discarded after resting the samples on a magnetic rack, and the beads were washed 3 times with 80% ethanol. The digestion was carried out by adding digestion solution (20 ng/ul trypsin in 50 mM HEPES, pH 8.5) to the beads (with protein bounded) at a 1:50 protease to protein ratio, at 37°C on a shaker overnight. The digested peptides were labeled with the TMTpro 18-plex kit (Thermo Fisher) according to the manufacturer protocol. All 18 channels were mixed equally and desalted before further analysis. The TMT labeled peptides were analyzed by an optimized two-dimensional liquid chromatography-tandem mass spectrometry (LC/LC-MS/MS) platform.

**Protein Identification and Quantification**

All the MS data were processed with an in-house JUMP software suite, a tag-based hybrid search engine to improve sensitivity. The protein database was generated by combining downloaded Swiss-Prot, TrEMBL, and UCSC databases and removing redundancy, followed by concatenation with a decoy database. Major parameters included 15 ppm mass tolerance for precursor ions and 20 ppm for product ions, full trypticity, static modification of the TMTpro tags (+304.20714 Da) on Lys residues and peptide N termini and carbamidomethyl modification on cysteine (+57.02146 Da), dynamic modification for Met oxidation (+15.99492 Da), maximal miscleavage sites (n=2), and maximal modification sites (n=3). The resulting PSMs are filtered to reduce protein FDR below 1%. Peptides generated from multiple homologous proteins were assigned to the canonical protein form in the manually curated Swiss-Prot database based on the rule of parsimony. If no canonical form was defined, the peptide was assigned to the protein with the highest PSM number. The protein quantification was based on TMT reporter ion intensities with $y_1$-ion based correction of TMT data to reduce the effect of ratio compression[13].

**Statistical Analysis**

Statistical analysis was carried out using GraphPad Prism version 9.0 (GraphPad Software, La Jolla, CA). Tests were performed only when experiments included three or more replicates. For comparing two groups, paired or unpaired two-tailed t tests were used. For experiments involving three or more groups, one-way or two-way ANOVA was applied, depending on the design. Details of the tests used are provided in the figure legends.

**Data availability**

The authors declare that the data supporting the findings of this study are available within the manuscript and its Supplementary Information. For material requests, contact the corresponding authors:. DNA sequencing and TMT data has been deposited to the Gene Expression Omnibus (GEO) under the accession no. GSE XXXXXX or to ProteomeXchange with accession no. PXDxxxxxx, respectively.

**References**


1. Watson, J. L. *et al.* De novo design of protein structure and function with RFdiffusion. *Nature* **620**, 1089–1100 (2023).
2. Chao, G. *et al.* Isolating and engineering human antibodies using yeast surface display. *Nat. Protoc.* **1**, 755–768 (2006).

3. Ghoda, L., Sidney, D., Macrae, M. & Coffino, P. Structural elements of ornithine decarboxylase required for intracellular degradation and polyamine-dependent regulation. *Mol. Cell. Biol.* **12**, 2178–2185 (1992).

4. Ng, A. H. *et al.* Modular and tunable biological feedback control using a de novo protein switch. *Nature* **572**, 265–269 (2019).

5. Lafleur, E. A. *et al.* Increased Fas Expression Reduces the Metastatic Potential of Human Osteosarcoma Cells. *Clin. Cancer Res.* **10**, 8114–8119 (2004).

6. Nguyen, P. *et al.* Route of 41BB/41BBL Costimulation Determines Effector Function of B7-H3-CAR.CD28ζ T Cells. *Mol. Ther. Oncolytics* **18**, 202–214 (2020).

7. Talbot, L. J. *et al.* Redirecting B7-H3.CAR T Cells to Chemokines Expressed in Osteosarcoma Enhances Homing and Antitumor Activity in Preclinical Models. *Clin. Cancer Res. Off. J. Am. Assoc. Cancer Res.* **30**, 4434–4449 (2024).

8. Prinzing, B. *et al.* Deleting DNMT3A in CAR T cells prevents exhaustion and enhances antitumor activity. *Sci. Transl. Med.* **13**, eabh0272 (2021).

9. Chen, Z. *et al.* The drug-elicitable alternative splicing module for tunable vector expression in the heart. *Nat. Cardiovasc. Res.* **4**, 938–955 (2025).

10. Abdelsamed, H. A. *et al.* Beta cell-specific CD8+ T cells maintain stem cell memory-associated epigenetic programs during type 1 diabetes. *Nat. Immunol.* **21**, 578–587 (2020).

11. Hudson, W. H. *et al.* Proliferating Transitory T Cells with an Effector-like Transcriptional Signature Emerge from PD-1+ Stem-like CD8+ T Cells during Chronic Infection. *Immunity* **51**, 1043-1058.e4 (2019).

12. Hughes, C. S. *et al.* Single-pot, solid-phase-enhanced sample preparation for proteomics experiments. *Nat. Protoc.* **14**, 68–85 (2019).

13. Wang, X. *et al.* JUMP: a tag-based database search tool for peptide identification with high sensitivity and accuracy. *Mol. Cell. Proteomics MCP* **13**, 3663–3673 (2014).
