## Supplemental Figures for "Reprogramming CAR T-Cells with designed bioPROTACs"

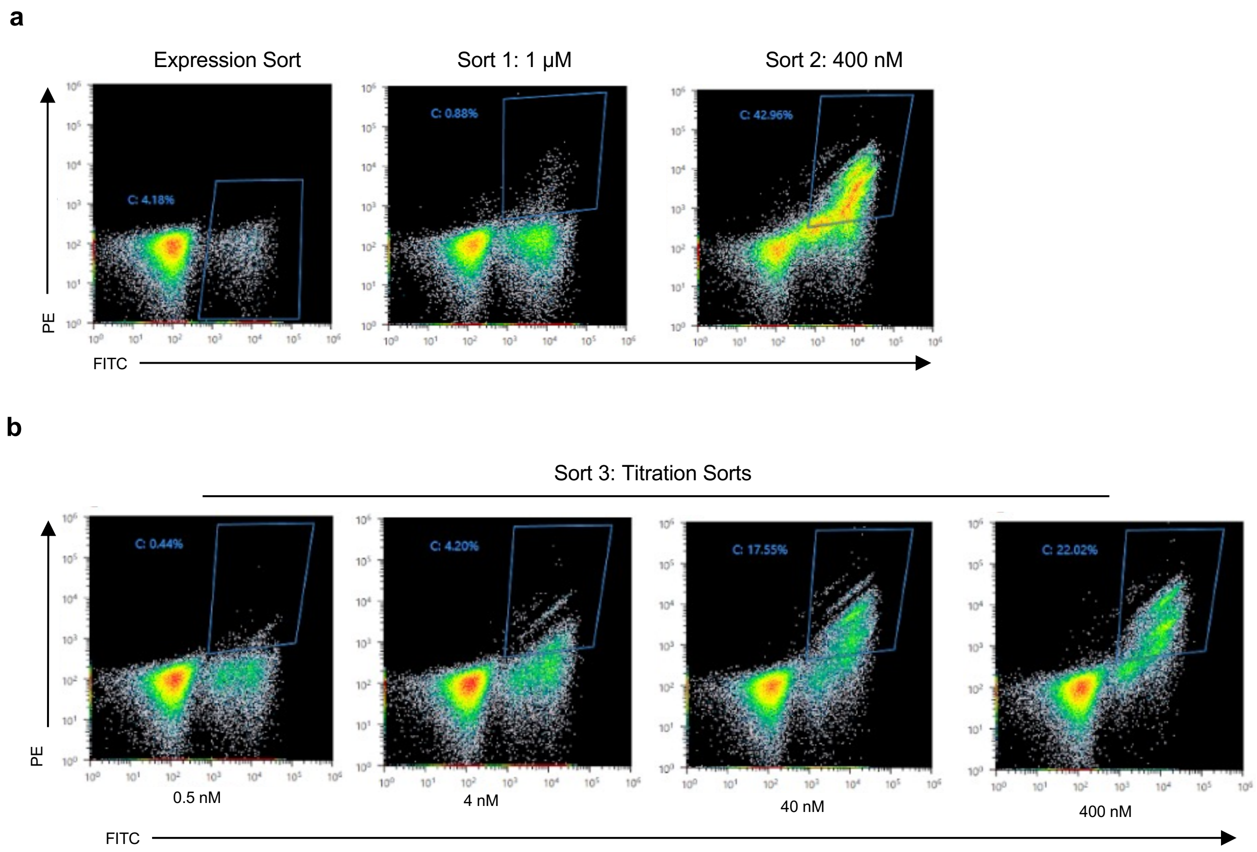


**Figure S1: Yeast surface display**

**a)** Initial sequential flow cytometry enrichment of yeast expressing DNMT3A binders, followed by addition of 1 μM and 400 nM of FLAG-tagged DNMT3A after subsequent enrichments.

**b)** Final round of flow cytometry titration sorts showing binder enrichment at 0.5 nM, 4 nM, 40 nM, and 400 nM of FLAG-tagged DNMT3A.

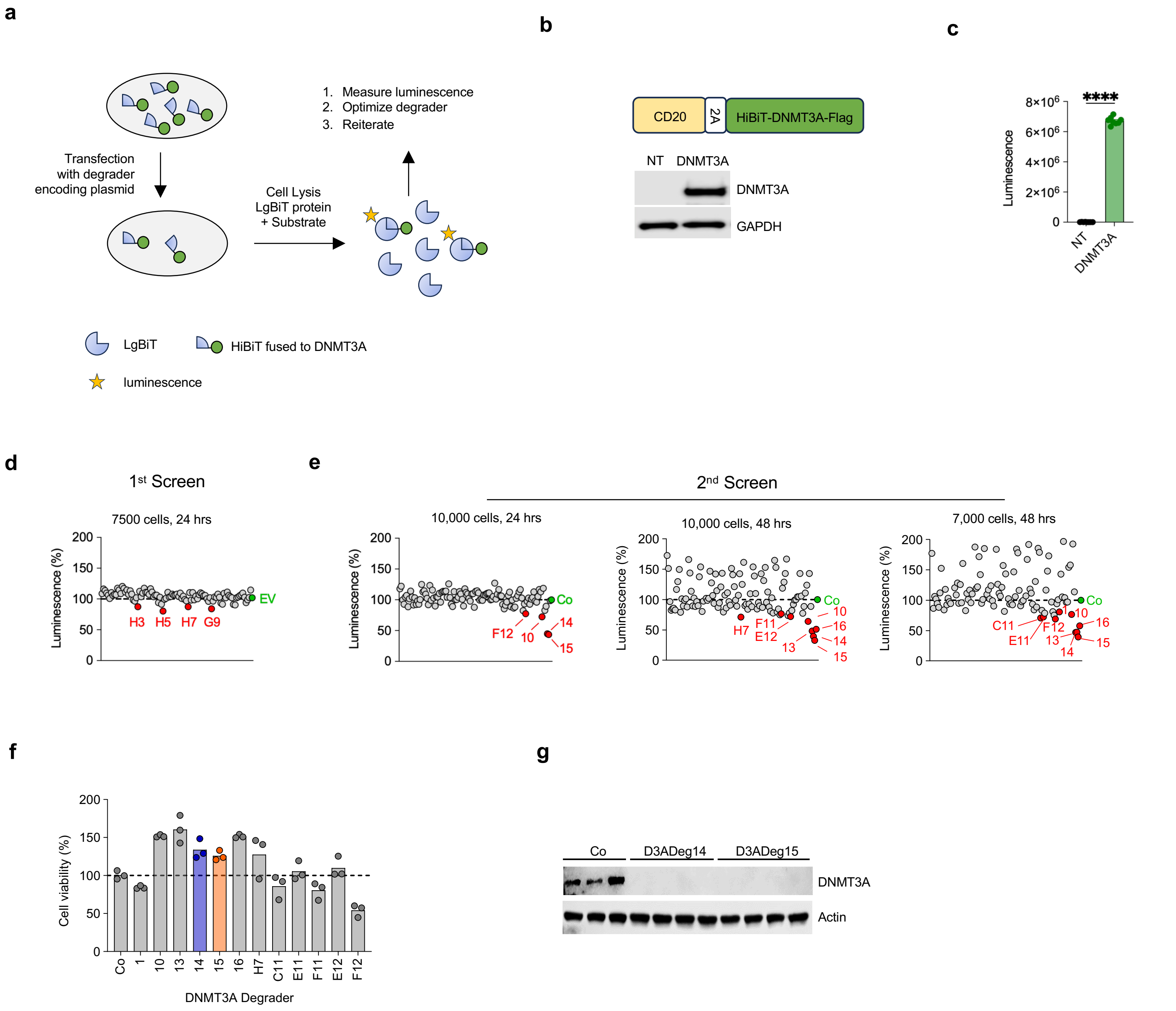


**Figure S2: DNMT3A bioPROTAC screen**

**a)** Screen schematic.

**b)** Retroviral construct encoding CD20 and HiBiT DNMT3A fusion. Western blot of non-transduced (NT) and transduced 293T cells (D3A 293T cells).

**c)** Bioluminescence assay comparing NT and transduced 293T cells.

**d)** Result of first screen; Co: control (empty vector).

**e)** Result of second screen. Screen was conducted at two cell concentrations.

**f)** Cell viability of potential hits shown in Figure 1c.

**g)** Detection of DNMT3A by Western blot in D3A 293T cells transfected with D3A.Deg14 and D3A.Deg15; Co: control (empty vector).

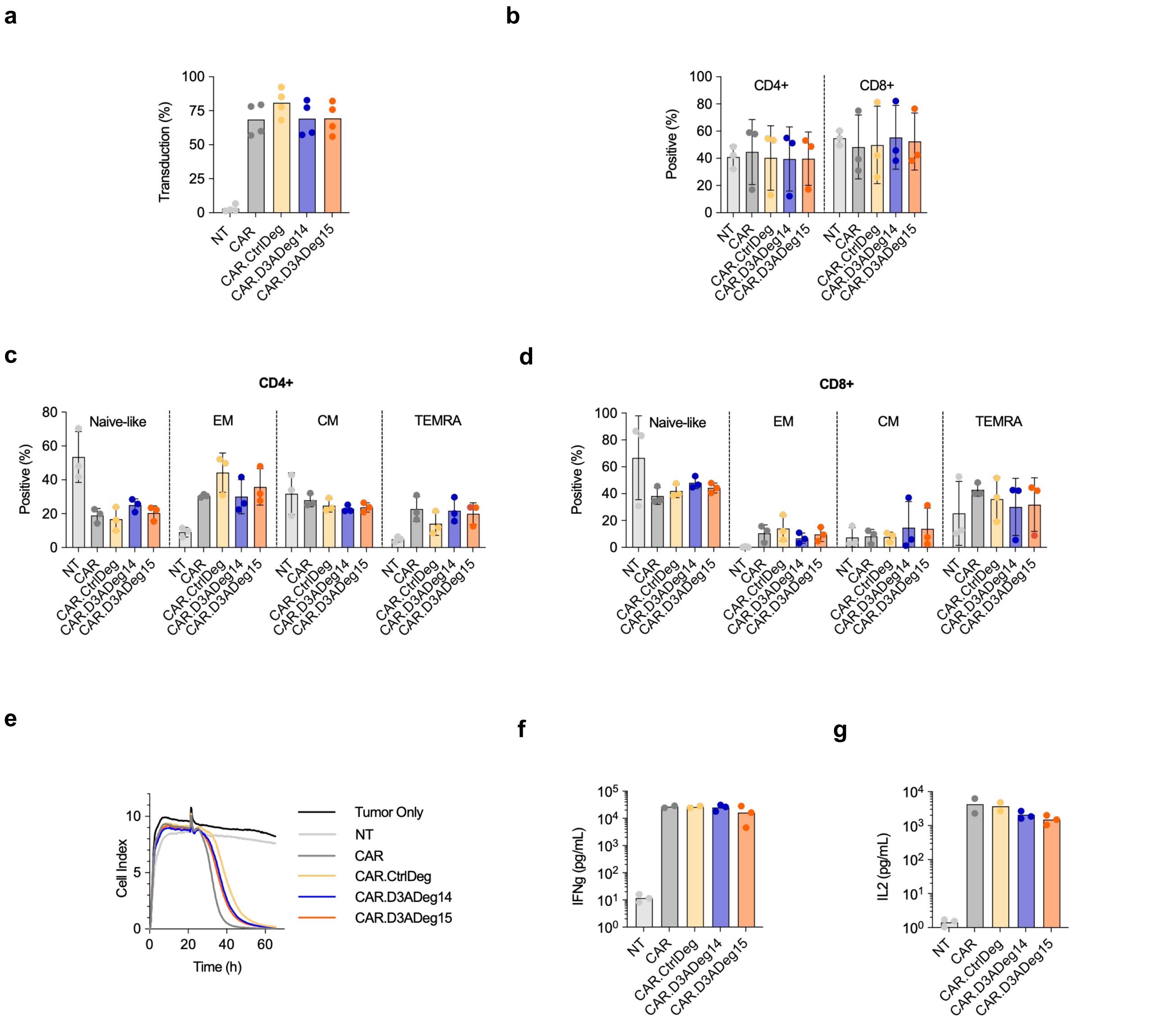


**Figure S3: Characterization of DNMT3A bioPROTAC expressing CAR T-cells**

**a)** Transduction efficiency at day 7 post-transduction assessed by flow cytometry for CAR (n=4)

**b-d)** Immunophenotype of CAR T cells assessed by flow cytometry; frequency of **(b)** CD4$^+$ and CD8$^+$ T cells. T cell differentiation state of **(c)** CD4$^+$ **(d)** and CD8$^+$ **(d)** T cells. Effector memory (EM): CCR7$^-$ CD45RA$^-$, central memory (CM: CCR7$^+$ CD45RA$^-$), naïve-like: CCR7$^+$ CD45RA$^+$, and T-effector memory re-expressing CD45RA (TEMRA): CCR7$^-$ CD45RA$^+$, (n=3).

**e)** Cytolytic activity of NT and indicated CAT T cell populations measured using xCELLigence real-time cell analysis (RTCA). B7-H3$^+$ LM7 tumor cells were co-cultured at an effector-to-target ratio of 2:1. One representative donor is shown, performed in triplicate.

**f-g)** NT and indicated CAR T cell populations were co-cultured with LM7 cells at a 2:1 E:T ratio. After 24 hours, **f)** IFNγ and **g)** IL-2 concentrations in the media were quantified by ELISA (n=2-3).

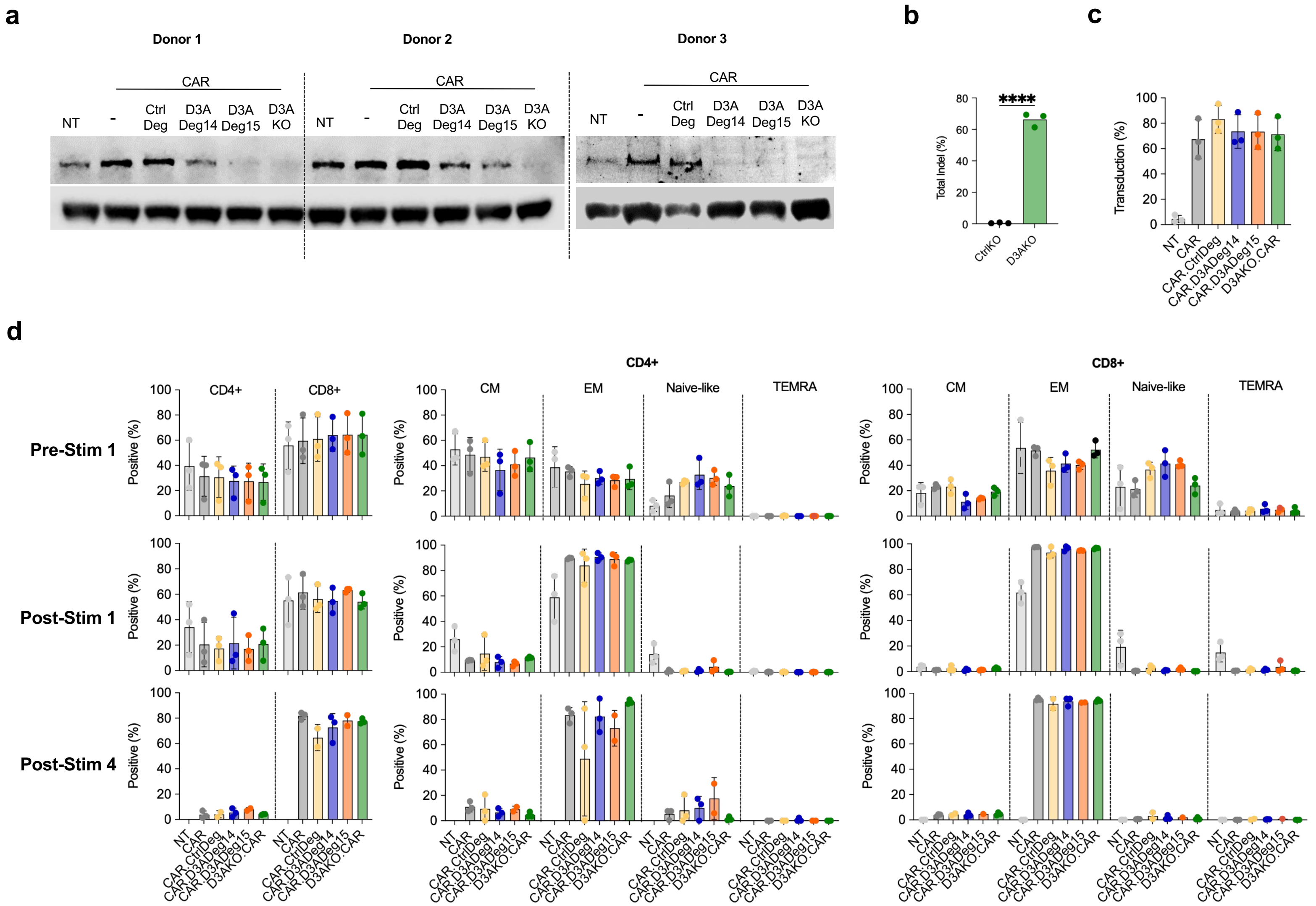


**Figure S4: Characterization of CAR T-cell populations**

**a)** Western blot confirming DNMT3A degradation (CAR.D3ADeg14, CAR.D3ADeg15) and deletion (D3AKO.CAR) on day 7 post-transduction (n=3).

**b)** INDEL analysis of DNMT3A KO T cells (****P < 0.0001, n=3).

**c)** Transduction efficiency of NT and indicated CAR T-cell populations control assessed on day 7 post-transduction by flow cytometry.

**d)** Immunophenotype of CAR T cells assessed by flow cytometry pre-stimulation (stim) 1, and after (post-stim) 1 and 4 stimulations (n=3).

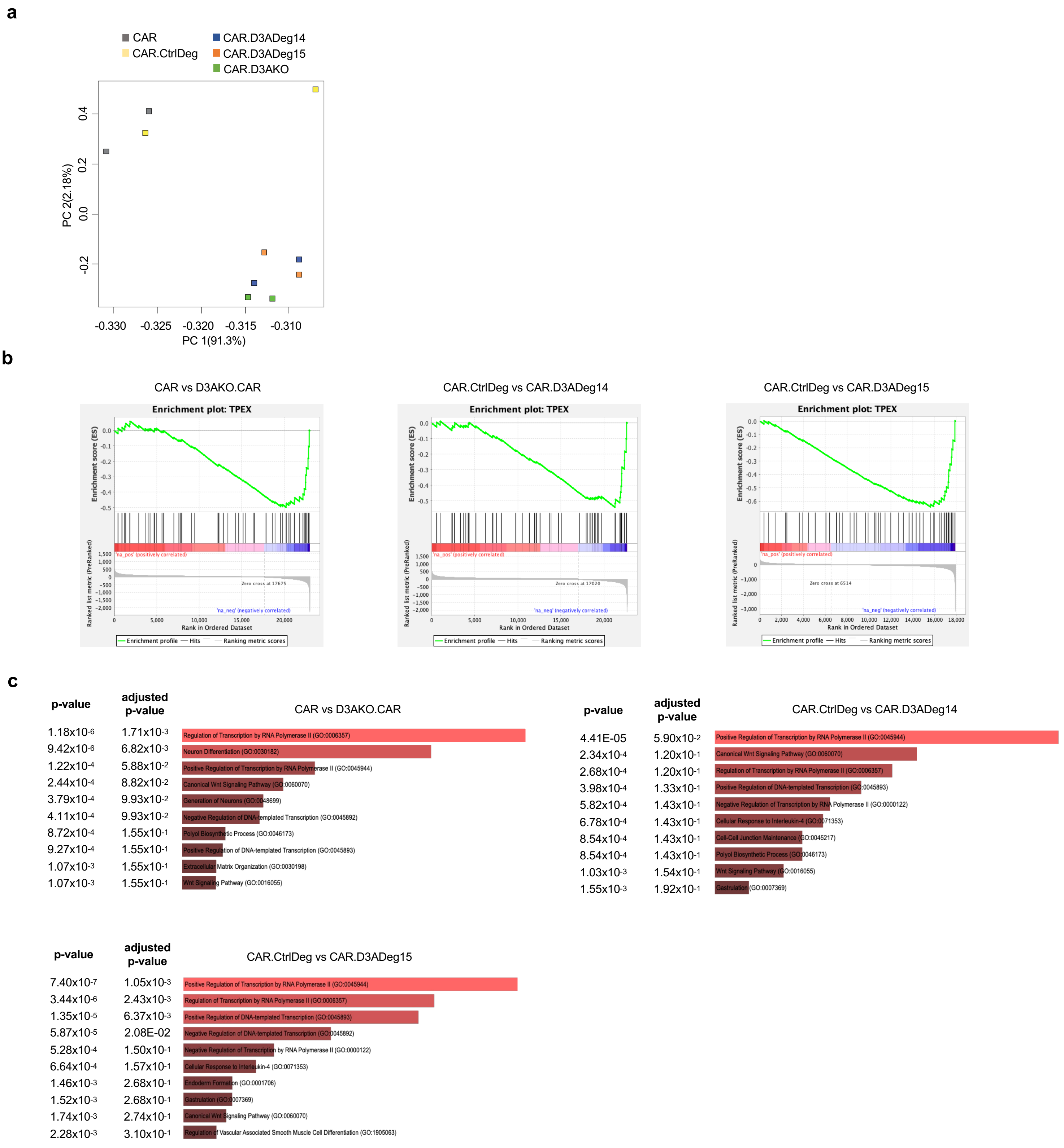


**Figure S5: DNMT3A bioPROTACs preserve CAR T-cell plasticity**

Figure S4 shows supplementary data analysis for the data shown in Figure 2a-h.

**a)** Principal Component Analysis (PCA) of DMR-based methylation profiles showing variance across CAR, CAR.CtrlDeg, CAR.D3ADeg14, CAR.D3ADeg15, and D3AKO.CAR conditions.

**b)** GSEA of DMRs after 4$^{th}$ repeat stimulation using a gene signature associated with progenitor exhausted T cells (TPEX). The following comparison are shown: CAR vs D3AKO.CAR, CAR.CtrlDeg vs CAR.D3ADeg14, and CAR.CtrlDeg vs CAR.D3ADeg15.

**c)** GO analysis of DNMT3A-regulated genes after 4$^{th}$ repeat stimulation. The following comparison are shown: CAR vs D3AKO.CAR, CAR.CtrlDeg vs CAR.D3ADeg14, and CAR.CtrlDeg vs CAR.D3ADeg15 with p-values, Fisher's exact test.
